# Advancing Cardiac Tissue Engineering: Melt Electrowriting Conductive Polymer-Hydrogel Scaffolds

**DOI:** 10.64898/2026.07.27.740898

**Authors:** M. Amini, J. Valdes-Fernandez, J. Latasa, E. Larequi, I. Anaut-Lusar, F. Prosper, M. M. Mazo Vega, A.M. Bittner

## Abstract

Myocardial infarction highlights an urgent need for strategies to regenerate functional cardiac tissue. Cardiac tissue engineering offers a promising approach; however, fabricating scaffolds that simultaneously integrate precise architectural anisotropy, mechanical compliance, and electrical conductivity remains an open challenge. In this work, we utilized melt electrowriting (MEW) to construct well-defined, 20-layer anisotropic rhomboidal polycaprolactone (PCL) scaffolds. We characterised them by tensile testing and by micro- and nanoscale microscopy. While introducing electrical conductivity via bulk blending with fillers (polypyrrole (PPy), polyaniline, or graphene oxide) compromised MEW print fidelity and failed to achieve physiological conductivity, surface coating strategies effectively combined conductivity from structural mechanics. Electrical and mechanical testing revealed that gold sputter coating and in situ PPy polymerization both imparted robust electrical conductivity while preserving the microfibrous architecture. However, when seeded with human induced pluripotent stem cell-derived cardiomyocytes (hiPSC-CMs) in fibrin hydrogels, only the gold-coated scaffolds supported synchronized, robust, and sustained contractile activity. PPy-coating resulted in functionally restricted constructs, suggesting that excessive structural rigidity limited tissue deformability. Gene expression analysis further revealed that elevated electrical conductivity alone does not drive hiPSC-CM maturation. Our data indicates that successful cardiac patch design relies on the integrated optimization of mechanics and architecture rather than treating conductivity as an isolated parameter, offering foundational guidelines for developing translational bioengineered heart patches.

**Graphical Abstract:** 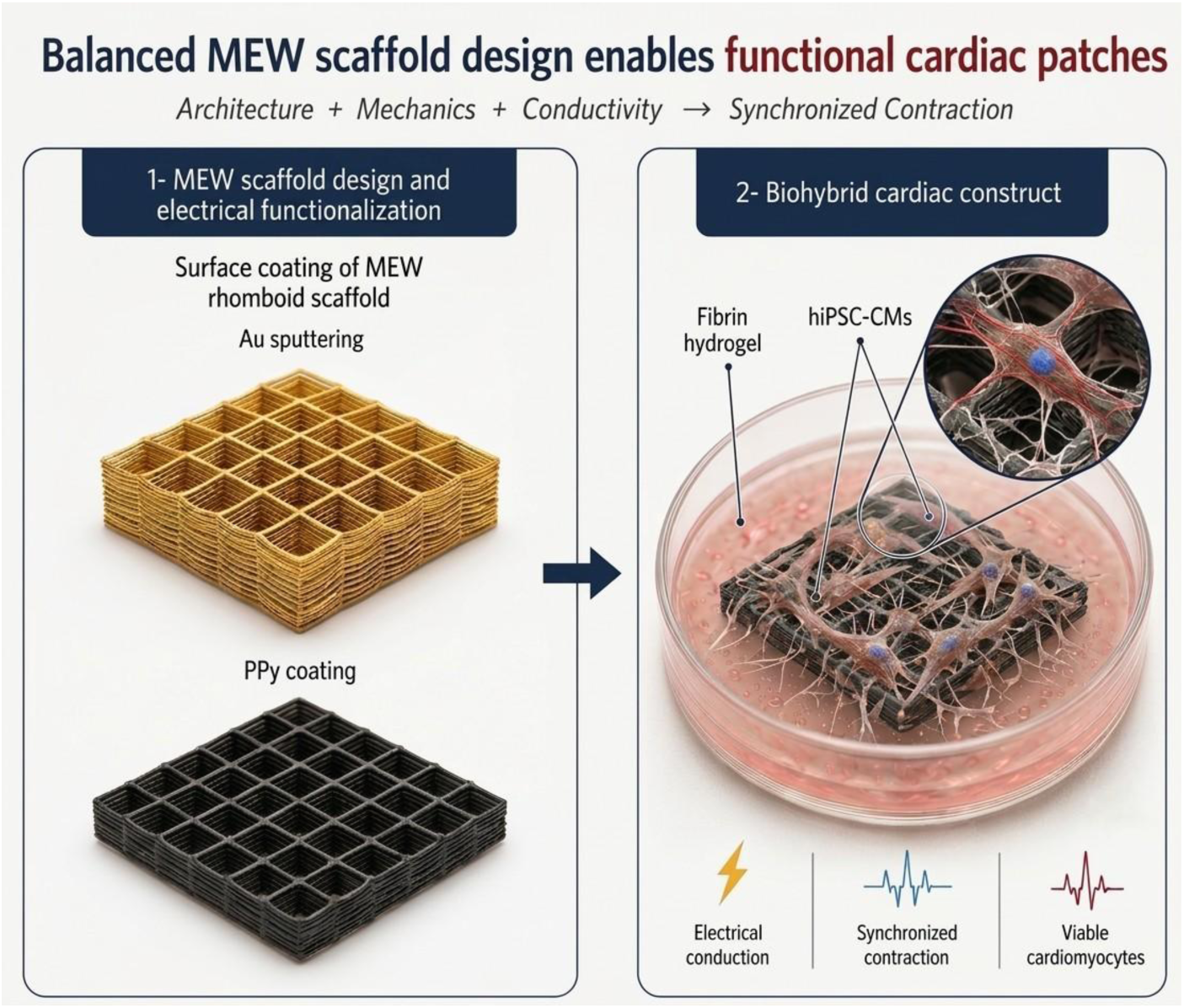

## 1. Introduction

Cardiovascular diseases (CVDs), including myocardial infarction (MI), remain the leading cause of mortality worldwide, posing major healthcare challenges [1]. MI results in the loss of an important fraction of cardiomyocytes (CMs), diminishing the population of force-producing units and disrupting tissue organization. This, together with the consequent fibre disarrangement and loss of electrical conductivity, compromise overall contractile performance. When patients survive an acute MI, the event often marks the onset of a chronic cardiovascular pathology, with progressive deterioration of cardiac function over time [2], [3]. Current pharmacological and interventional strategies are essentially palliative, aiming to alleviate symptoms and delay disease progression, while heart transplantation remains the only truly therapeutic option. However, transplantation is constrained by donor organ shortages and risks of immune rejection [4], [5]. Thus, all of the above, together with the incapacity of human myocardium to regenerate [6], underscore the urgent need to generate human cardiac tissue in the laboratory as a foundation for novel therapeutic approaches. Cardiac tissue engineering (CTE) has emerged as a promising strategy by aiming to biofabricate damaged heart tissue from engineered constructs that closely replicate the structural and functional properties of native myocardium [7].

Tissue engineering strategies pivot on two fundamental components: a cellular source that provides the biological functionality of the generated constructs, and a scaffold that supports the cells and enables them to perform their function [8]. The careful selection of both elements is critical for the development of successful therapeutic approaches, and the simultaneous advances in both cellular reprogramming and additive manufacture strategies provide an ideal opportunity to significantly improve CVD treatment and patient life quality [9].

Regarding the cellular component, the advent of human induced pluripotent stem cell (hiPSC) technology has revolutionized the field by enabling access to virtually any cardiac lineage [10]. The development of chemically defined, monolayer-based differentiation protocols has become the gold standard due to their efficiency, stability, and reproducibility [11]. HiPSC-derived CMs (hiPSC-CMs) are being extensively investigated for MI therapy. Early strategies involving intramyocardial injection showed limited long-term benefits due to poor survival, arrhythmogenic risk, and scalability issues, despite some encouraging outcomes [12]. More advanced approaches, such as cardiac patches combining hiPSC-CMs with biomaterials, have demonstrated improved repair in animal MI models [13]. Finally, combining hiPSC-CMs with rhomboidal PCL scaffolds, i.e. with scaffolds based on a rhombus-shaped unit cell, has been shown to enhance cardiac function in rat MI models by promoting cellular alignment and integration into native myocardium [14].

In parallel with cellular advances, the scaffold component has become equally critical for successful CTE. Beyond simply providing structural support, scaffolds must recreate the physical and biochemical cues that guide CM function. Their composition, mechanical stiffness, and electrical properties influence not only cell survival and phenotype, but also the synchronization of contraction and long-term tissue integration. Thus, designing scaffolds that combine adequate mechanical strength with physiological compliance and conductivity is fundamental for reproducing the native myocardial environment and enabling functional tissue regeneration [15]. To achieve this, several considerations must be considered.

Material selection plays a pivotal role in scaffold design. In CTE, natural matrices such as collagen and fibrin are often considered the biological gold standards due to their native bioactivity, while poly(ε-caprolactone) (PCL) is widely used as a synthetic alternative because of its biocompatibility, biodegradability and excellent processability in additive manufacturing.

Precision in biofabrication is essential to accurately mimic the complex architecture of cardiac tissue. Traditional 3D printing methods often fall short in terms of resolution and precision required for fabricating such complex structures at the micron scale, resulting in suboptimal tissue constructs [7]. Despite fused deposition modelling (FDM) generates 3D scaffolds with good mechanical integrity, the millimetre-scale fibres produced hinder integration with soft, cell-laden hydrogels, reducing biological performance [11]. To overcome these limitations, Melt Electrowriting (MEW) has emerged as a powerful approach to fabricate microfibrous scaffolds with diameters between 10 and 100 µm. While native extracellular matrix (ECM) structural proteins primarily possess nanoscale diameters, the microscale topography provided by MEW closely matches the cellular dimensions of cardiomyocytes, thereby serving as an effective biomimetic architectural template that facilitates contact guidance, robust attachment, and cellular alignment [8,9]. MEW-printing strategies are continuously optimized to create scaffolds with finer fibres that better mimic the ECM, though their reliance on nonconductive PCL restricts functional integration in cardiac applications [12].

Scaffold geometry and fibre orientation are crucial in guiding CM alignment, influencing mechanical performance and biological outcomes [4]. Scaffold geometry optimization studies demonstrated that stacked rhomboid patterns maximize cardiomyocyte alignment and engineered tissue performance [13]. Yet, to the best of our knowledge, systematic optimization of the rhomboid design in terms of unit cell size or the number of layers, both critical parameters influencing mechanical anisotropy and biological performance [5,7], has not been reported.

Beyond mechanical and structural fidelity, electrical conductivity is important for engineered cardiac tissues, supporting synchronized contraction and functional integration of the construct with the host myocardium [16]. Conductive scaffolds facilitate the propagation of electrical signals between CMs across the engineered tissue, thereby strengthening electromechanical coupling [17]. When combined with non-conductive fibrin hydrogels, conductive scaffolds can form engineered cardiac patches and restore both mechanical and electrical function following MI [12]. The requirement of stretchability and elasticity suggests using again polymers in or on the MEW fibres, most simply well-known organic semiconductors, which require substantial levels of doping. The performance of such scaffolds has been addressed before [12,17], but to our knowledge never under CM contraction, although the combined electrical and mechanical behaviour is key to high performance.

Hence, we have developed and systematically optimized electrically conductive PCL-MEW scaffolds with rhomboid geometries, specifically tailored to improve hydrogel integration. We have optimized MEW printing parameters, scaffold structure and unit cell size, to promote cardiomyocyte alignment, improve biological performance, and overcome the electrically isolating nature of PCL. To better mimic cardiac tissue conductivity, two different strategies were evaluated, bionanomaterial integration and surface coating, respectively. A thorough morphological, mechanical, physicochemical and electrical characterization was carried out. We then assessed the efficacy of the conductive scaffolds in an in vitro model in which they were combined with hiPSC-CMs, with the construct embedded in a fibrin hydrogel. Naïve PCL scaffolds were employed as controls to evaluate whether the incorporation of conductive elements enhanced tissue performance in terms of CM survival, contractile activity, and tissue maturation.

## 2. Materials and methods

### 2.1. Melt Electrowriting

Highly organized microfibrous scaffolds were fabricated using a custom-built MEW system (NovaSpider, [18]). The setup consisted of a static collector and a nozzle capable of precise movement along the X, Y, and Z axes, where the XY plane defined the printing area and the Z-axis controlled the nozzle-to-collector distance. PCL (medical grade PCL, Sigma-Aldrich) was chosen due to its MEW-compatible thermal properties and medical-grade availability. All the tests were done with a nozzle diameter of 4 mm, at room temperature, and a relative humidity of 50 ± 5%.

### 2.2. Fabrication of Conductive Polymer Scaffolds

Conductive scaffolds were produced either by blending conductive fillers with PCL using a solution mixing method or by modifying the surface of PCL scaffolds through coating techniques. The conductive materials incorporated in this study were polypyrrole (PPy, 98%, Sigma-Aldrich), polyaniline (PANI, emeraldine base, ≥98%, Sigma-Aldrich), and graphene oxide (GO dispersion in water, 4 mg/mL, Graphenea)

#### 2.2.1. Preparation of Conductive Fillers

PPy was used as received. PANI was converted from the emeraldine base to its conductive salt form. For this purpose, HCl solutions (10, 20, and 30% w/w, Sigma-Aldrich) were prepared in ultrapure water (Milli-Q® Advantage A10, Merck Millipore, resistivity 18 MΩ·cm, TOC < 10 ppb), and emeraldine base powder was added under gentle stirring to ensure complete protonation. The emeraldine salt was then isolated by filtration, washed with water to remove excess acid, and dried at room temperature overnight. For GO, a dispersion in water (250 μL containing 1 mg GO) was centrifuged at 2000 rpm (rotor radius = 15 cm), redispersed in 1 mL acetone, and used for subsequent mixing with PCL.

#### 2.2.2. Solution Mixing Method

For the PCL–PPy blend, 1.47 g of PCL was dissolved in 12 mL DMF, while 0.03 g of PPy, corresponding to 2 wt.% relative to the total polymer content, was dispersed in 12 mL DMF. The two solutions were mixed, stirred at room temperature, and sonicated at 50 °C for 2 h. The resulting blend was dried at ambient temperature overnight.

The PCL–PANI blend was obtained using the same DMF-based solution mixing method as described above for the PCL–PPy system. A concentration of 5 wt.% PANI salt prepared using 10% HCl was incorporated into PCL for this blend.

The PCL–GO nanocomposite was synthesized by dissolving 1 g of PCL in 10 mL DMF and adding the redispersed GO (1 mg GO in 1 mL acetone). The mixture was sonicated for 1 h and stirred for 30 min to promote particle distribution. Afterwards, the PCL–GO mixture was dried to remove solvent, and subsequently melted for MEW processing.

#### 2.2.3. Surface Coating Methods

Conductive surface modification was carried out via in situ polymerization of pyrrole on the surface of PCL scaffolds. The scaffolds were immersed in an aqueous pyrrole solution, followed by the addition of FeCl₃ as the oxidizing agent and p-toluenesulfonic acid (pTS) as the catalyst. Three coating conditions were investigated: (i) a 6 h reaction using 0.1 M pyrrole, 0.27 M FeCl₃, and 0.1 M pTS; (ii) a 12 h reaction using 0.1 M pyrrole, 0.27 M FeCl₃, and 0.1 M pTS; and (iii) a 6 h reaction using 0.05 M pyrrole, 0.135 M FeCl₃, and 0.05 M pTS. All solutions were prepared in Milli-Q water, and the polymerization was carried out under identical conditions.

Gold was deposited on PCL scaffolds by sputter coating in a Quorum Q150T ES system. The chamber was evacuated to ∼ 10⁻⁴ mbar and argon gas introduced to 10⁻² mbar. Gold deposition was performed at 20 mA for ∼ 15 min, resulting in a uniform conductive film. The actual coating thickness was then measured by X-ray reflectivity (XRR) using an X’Pert PRO PANalytical X-ray diffractometer equipped with a PIXcel 3D detector. The analysis revealed that the actual gold layer thickness was ∼ 10% higher than the nominal value set by the sputtering system.

### 2.3. Characterization

#### 2.3.1. Structural and chemical characterization

Raman and Fourier-transform infrared (FTIR) spectroscopy were recorded at ambient conditions. The Raman tool was an Alpha300 R Raman microscope (WITec) equipped with a 532 nm laser excitation source. The spectra were recorded in the range of 100–4000 cm⁻¹ with a resolution of 3 cm⁻¹. They were smoothed with a 3-point averaging filter, followed by baseline extraction and subtraction. Multiple spots on each sample were measured to ensure signal reproducibility and surface homogeneity. FTIR spectroscopy was carried out in a PerkinElmer Frontier FT-IR spectrometer in attenuated total reflectance (ATR) mode. Samples were pressed against a diamond ATR prism, referenced against air. The resolution was 4 cm⁻¹, and spectra were averaged over 100 scans and smoothed with a 6-point averaging filter.

#### 2.3.2. Morphological and surface characterization

The surface morphology and topographical features of the fibres and fabricated scaffolds were analysed using environmental scanning electron microscopy (ESEM) and atomic force microscopy (AFM). ESEM imaging was performed in a FEI Quanta 250 microscope (Thermo Fisher) operated in low vacuum mode (water vapor pressure 10-200 Pa). Uncoated samples were fixed on aluminium stubs with conductive carbon tape. Images were usually recorded with an accelerating voltage of 10 kV. This enabled detailed visualization of fibre alignment, diameter, and scaffold porosity. The AFM tool was a JPK NanoWizard V (Bruker) in AC mode under ambient conditions. Multi 75 Silicon cantilevers (budgetsensors) with a nominal tip radius of <10 nm were used. The scan range was set between 1 and 5 µm, to capture representative topographical features. Simultaneously acquired images of the phase of the AC signal were used to highlight height variations.

#### 2.3.3. Electrical characterization

Two complementary methods were employed to assess the electrical conductivity of the scaffolds, depending on the type of sample. AC Broadband Dielectric Spectroscopy (BDS) was used to evaluate the bulk conductivity of PCL composites containing conductive fillers (PPy, PANI, or GO) prepared via solution mixing. In contrast, surface conductivity of the scaffolds coated with gold or PPy was measured with the DC four-point probe technique. All experiments were carried out under ambient conditions.

Bulk complex dielectric conductivity was measured using a Novocontrol Alpha-S analyser over a frequency range of 10⁻¹ to 10⁶ Hz at 37 °C, utilizing a fixed AC voltage amplitude of 1.0 V. For powder samples, a capacitor configuration was assembled using a DSC pan and a 4 mm parallel-plate electrode. To prevent short-circuiting, a fluoropolymer film was inserted between the electrode and the wall of the pan. The sample was placed in the pan and gently compacted by applying a uniaxial pressure of 5–10 MPa for a few seconds to ensure good particle contact and minimize porosity. The top electrode was then positioned to complete the measurement setup.

Surface conductivity measurements were carried out in a four-point probe station (Jandel Engineering Ltd.). The setup included tungsten-carbide probe tips, typically 0.4 mm in diameter with ∼100 µm radius, and a Jandel RM3000(+) test unit, serving as both the current source and voltmeter. Scaffolds were placed on flat, non-conductive substrates to avoid interference. The electrical properties were evaluated directly via the measured sheet resistance (R_s_, in Ω/sq) under varying current biases. This approach allowed for a thickness-independent comparative analysis of the electrical pathways formed by the thin surface functionalization layers (gold and PPy) across the microfibrous meshes. This method provides highly accurate and reproducible electrical profiling and is particularly well-suited for evaluating thin surface coatings.

#### 2.3.4. Mechanical characterization

##### 2.3.4.1. Uniaxial tensile tests

Uniaxial tensile tests were performed using a CERT UMT testing machine, for straight fibres and scaffolds. Straight fibres, printed in 3, 5, and 10 layers and tested with a gauge length of 6 mm, were stretched at a strain rate of 2 mm/s until failure to assess their intrinsic strength and stretchability. Rhomboid-patterned scaffolds were printed with 10 and 20 layers; each scaffold was composed of repeating rhomboid-shaped unit cells, subdivided into either 60 or 120 segments per side, resulting in arrays of 60×60 and 120×120-unit cells. Three specific configurations were examined in detail: 10 layers with 60×60 rhomboid unit cells (10-layer R60), 20 layers with 60×60-unit cells (20-layer R60), and 20 layers with 120×120-unit cells (20-layer R120). In R120, the short and long diagonals measured 0.5 mm and 1 mm, respectively, while in R60, the short diagonal was 1 mm and the long diagonal was 2 mm. Scaffold specimens with a gauge length of 20 mm were stretched at a strain rate of 2 mm/s (10%/s) until failure. This experimental setup allowed a comprehensive comparison of how scaffold geometry, layer count, and unit cell size influence tensile strength, stiffness, and failure mechanisms.

##### 2.3.4.2. Cyclic tensile tests

To replicate cardiac cycling behaviour, cyclic tensile loading was applied to 20-layer R60 and R120 scaffolds at 5% and 10% strain. The 5% strain corresponds to an elongation from 20 mm to 21 mm. All tests were conducted under fixed conditions, namely a total of 20 loading–unloading cycles, applied at a strain rate of 0.4 mm/s (for 20 mm samples this corresponds to 0.02/s). The cyclic mechanical response was recorded by plotting load–displacement curves.

### 2.4. hiPSC culture and differentiation

All experiments were carried out using the WTC11 hiPSC line (UCSFi001-A), generously provided by Prof. Bruce Conklin (Gladstone Institutes). Cells were maintained in E8 medium on plastic surfaces coated with growth factor–reduced Matrigel (GFR-MG, 1:180 dilution) and routinely passaged at a 1:15 ratio using 0.5 mM EDTA (Invitrogen) every 4–5 days. Cardiac differentiation was induced by a biphasic modulation of the Wnt signalling pathway, following the protocol established by the Burridge laboratory [19]. Once hiPSCs reached confluence, the culture medium was switched to RPMI supplemented with B27 minus insulin (RPMI B27−) and 8 µM of the GSK3 inhibitor CHIR99021 (CHIR, Axon Medchem) for 24 h. Following this initial exposure, the concentration of CHIR was reduced to 3 µM and maintained for an additional 48 h. After CHIR supplementation, the medium was replaced with RPMI B27− supplemented with 5 µM of the Wnt inhibitor C59 (Axon Medchem) for 48 h. Cultures were then maintained in RPMI supplemented with full B27 (RPMI B27). By days 9–10, spontaneously beating CMs were observed. To enrich the CM population, cells underwent two consecutive rounds of metabolic selection by incubation for 72 h in glucose-free RPMI supplemented with 4 mM lactate. Between these two rounds, cells were dissociated with TrypLE (Gibco; 7–10 min at 37 ° C), carefully collected, and replated on GFR-MG–coated plates (1:80 in RPMI B27). The replating medium was supplemented with 10% Knock-Out Serum Replacement (KSR, Gibco) and 10 µM ROCK inhibitor Y-27632 (Y27, Tocris) to improve post-dissociation survival. After 24 h, the medium was replaced with RPMI B27 without Y27, and cultures were subsequently maintained in RPMI B27 with regular medium changes until use. The entire process required 22 to 25 days.

### 2.6. Cell isolation and tissue generation

Before seeding, PCL, Au and PPy scaffolds were punched into circular samples with a diameter of 6 mm and exposed to a 1 mbar O₂ plasma (Diener Electronic) for 5 minutes to enhance hydrophilicity and promote cell suspension penetration. The scaffolds were subsequently sterilized by incubation in 70% ethanol for 30 minutes, rinsed three times with sterile distilled water, and air-dried. At the end of the differentiation process, hiPSC-CMs were washed twice with 0.5 mM EDTA in PBS and dissociated into single cells using TrypLE (Gibco) for 7–10 minutes at 37 ° C. Cells were gently collected, pooled, and counted. Final cell density was adjusted to 7 × 10⁴ cells per mm²of scaffold. To prepare the tissue mastermix, the cell suspension was combined with RPMI B27 medium supplemented with 10% KSR, 1% penicillin/streptomycin, 10 µM Y-27632 (Y27), and 6 mg/mL bovine fibrinogen (Sigma Aldrich). For construct assembly, 35 µL of this mastermix together with 1.8 µL of thrombin (100 U/mL, Biopur) were dispensed into a Teflon mold containing the mPCL scaffolds, followed by incubation for 1 h at 37 ° C to allow fibrin polymerization. The constructs were then transferred to culture plates with RPMI B27 supplemented with 10% KSR, 1% penicillin/streptomycin, 10 µM Y27, and aprotinin (0.1% wt/vol; 33 μg/mL, Sigma Aldrich). After 24 h, the medium was replaced with RPMI B27 lacking Y27 but containing aprotinin to prevent fibrin degradation. Medium was subsequently renewed every other day.

### 2.7. Alamar Blue assay

Cellular metabolic activity was evaluated using the AlamarBlue viability assay (Invitrogen), according to the manufacturer’s protocol. Phenol red–free RPMI medium (Life Technologies) was supplemented with AlamarBlue to a final concentration of 10% (v/v), and the constructs were incubated in this solution for 2 h 30 min at 37 ° C. Afterwards, 100 µL of the conditioned medium was transferred to 96-well plates, and absorbance was quantified using a plate reader (BMG Labtech).

### 2.8. Gene expression analysis by qRT-PCR

Total RNA was extracted from differentiated cells and engineered cardiac tissues using Trizol reagent (Life Technologies), following the manufacturer’s guidelines. 1 µg of RNA was reverse-transcribed into cDNA with the PrimeScript RT Reagent Kit (Takara). Quantitative real-time PCR was performed using 5 ng of cDNA as template in PowerUp™ SYBR™ Green Master Mix (Applied Biosystems) on a QuantStudio 5 Fast Real-Time PCR system (Applied Biosystems). GAPDH served as the internal reference gene, and relative expression levels were calculated using the 2^−ΔΔCt^ method.

### 2.9. Statistical analysis

Data normality was assessed using the Shapiro–Wilk and Kolmogorov–Smirnov tests. Results are presented as mean ± SEM for normally distributed datasets or as median (Q1, Q3) for non-parametric data. Comparisons between two groups were carried out with Student’s t-test, while experiments involving more than two groups were evaluated using one-way or two-way ANOVA followed by Tukey’s HSD post hoc test. Statistical analyses were conducted with GraphPad Prism 8 (GraphPad Software Inc.), and differences were considered statistically significant at p < 0.05.

## 3. Results and discussion

### 3.1. Optimisation of MEW printing parameters

Optimizing MEW parameters is essential due to the high sensitivity of the process to ambient conditions, material properties, and desired scaffold geometries [20]. To establish a reproducible fabrication protocol, the key parameters governing fibre formation must be systematically evaluated. Of the parameters that control the jetting process, four have the most significant effect on fibre deposition and diameter: Voltage, pressure, translation speed and heating temperature. We will discuss their systematic optimisation here because we believe that it was essential to map out the exact processing window for our specific polymer-tool configuration, and to aligned it with established results.

While changes in voltage did not significantly affect the fibre diameter, printing was unsuccessful at 0 kV and inconsistent at low absolute voltages (< 4 kV), with a nozzle-to-collector distance of 5 mm, which is in accordance with Castilho et al. [20]. The median diameter increased from 14.3 µm (IQR: 9–15 µm) at −4 kV to 27.43 µm (IQR: 25–28 µm) at −7 kV (Figure S1A). Between −6 kV and −7 kV, fibres formed consistently. Interestingly, at −8 kV fibres remained consistent, but in ∼5% of cases, the high voltage caused the fibre to split into two parts: a thick, straight fibre and a thin, curly fibre (Figure S2A). This suggests that voltage primarily governs the jet initiation and stability rather than directly controlling fibre diameter, unfolding in agreement with the foundational principles established by Brown, Dalton, and Hutmacher [5].

Pressure changes have a significant impact on fibre diameter. The median fibre diameter more than quadruples from 8.82 µm (IQR: 5–9 µm) at 20 mbar to 37.70 µm (IQR: 32–45 µm) at 500 mbar (Figure S1B). At pressures below 20 mbar, particularly at 0 mbar, the fibres become noticeably thin and curly, with diameters sometimes reducing to several hundred nanometres (Figure S2B, C). Similar fibre curling has been reported by Castilho et al. [20] and Böhm et al. [21] when decreasing the translation speed; however, such behaviour has not previously been reported as a function of reduced pressure. The resulting disordered geometry and irregular fibre arrangement closely resemble the morphological characteristics of conventional melt electrospun meshes. This suggests that at near-zero pressures, the drastically reduced mass flow rate disrupts the stable jet deposition typical of MEW, leading to a printing regime that phenotypically mimics melt electrospinning.

By increasing the translation speed, the median fibre diameter is reduced from 30.58 µm (IQR: 29–32 µm) at a translation speed of 500 mm/min to 14.58 µm (IQR: 14–15 µm) at 4000 mm/min, less than half (Figure S2C). This phenomenon is again very well known [20, 21].

PCL cannot be printed below 70 °C, as the polymer becomes too viscous to be extruded in a stable, continuous manner. On the other end of the temperature range, when the polymer is heated above 150 °C, the behaviour of the jet changes dramatically, shifting from straight-line deposition to a sinusoidal or buckled pattern (Figure S2D). The reason of this thermally induced buckled pattern, which was to our knowledge not reported before, might be effects of reduced viscosity and increased flow rate, which compromise the positional stability of the jet. It is obvious that this affects the fibre diameter, too. We found that an increase in temperature from 70 °C to 120 °C led to a 56% increase in the median fibre diameter (Figure S1D). This increase is due to the lower viscosity at elevated temperatures, which allows for a greater material flow through the nozzle, leading to thicker fibres.

We found that the optimal balance of the four parameters is achieved at a voltage of −6 kV, an air pressure of 100 mbar, a nozzle temperature of 80 °C, and a translation speed of 800 mm min⁻¹ (for our usual nozzle-to-collector distance of 5 mm). These specific parameters were selected as optimal because they yielded the highest structural fidelity, characterized by continuous jet stability, strict straight-line deposition tracking, and zero fibre curling or pulsing. Under these optimized conditions, the scaffolds exhibited a highly reproducible median fibre diameter of 18 µm (IQR: 15–22 µm). These conditions are consistent with previously reported MEW studies. For instance, Iglesias-García et al. reported optimal parameters of 2 bar pressure, 6.5 kV voltage, 800 mm min⁻¹ translation speed, an 8 mm nozzle-to-collector distance, and a 23 G needle (≈330 µm ID), resulting in fibres with diameters of ∼ 17 µm [15]. Fuchs et al. employed a temperature of 73 °C, a 22 G nozzle, 1.2 bar pressure, a voltage of 6 kV, a 4 mm working distance, and a translation speed of 400 mm min⁻¹, yielding fibres of ∼20 µm diameter [22]. Saiz et al. reported optimal conditions including a 24 G nozzle, a 3.5 mm collector distance, a high voltage of 6.3 kV, a melt temperature of 80 °C, a pneumatic pressure of 2.4 bar, and translation speeds ranging from 260 to 460 mm min⁻¹ for different inks, which resulted in an average fibre diameter of 20.7 ± 0.9 µm [23].

### 3.2. Effect of layer number: Uniaxial tensile testing of multilayer MEW fibres

Uniaxial tensile testing of the MEW-fabricated PCL scaffolds revealed exceptional stretchability, which is crucial for cardiovascular surgery applications. As demonstrated in Figure 1, the scaffolds exhibited excellent maximum elongation values exceeding 800%, even for the 10-layer configurations. This remarkable stretchability before rupture far surpasses the typical mechanical behavior of bulk PCL, confirming the structural advantages of the microfibrous architecture. For comparison, previous studies on FDM–printed PCL have reported maximum elongations of approximately 120% [24]. The stress– strain curves exhibited a series of discrete stress drops, corresponding to the sequential rupture of individual fibre layers, and potentially also to interlayer sliding or partial layer detachment within the multilayer constructs. This progressive failure mode reflects the layered architecture of MEW constructs, which enables fibre sliding, reorientation, and gradual alignment under tensile loading rather than catastrophic fracture.

**Figure 1.**
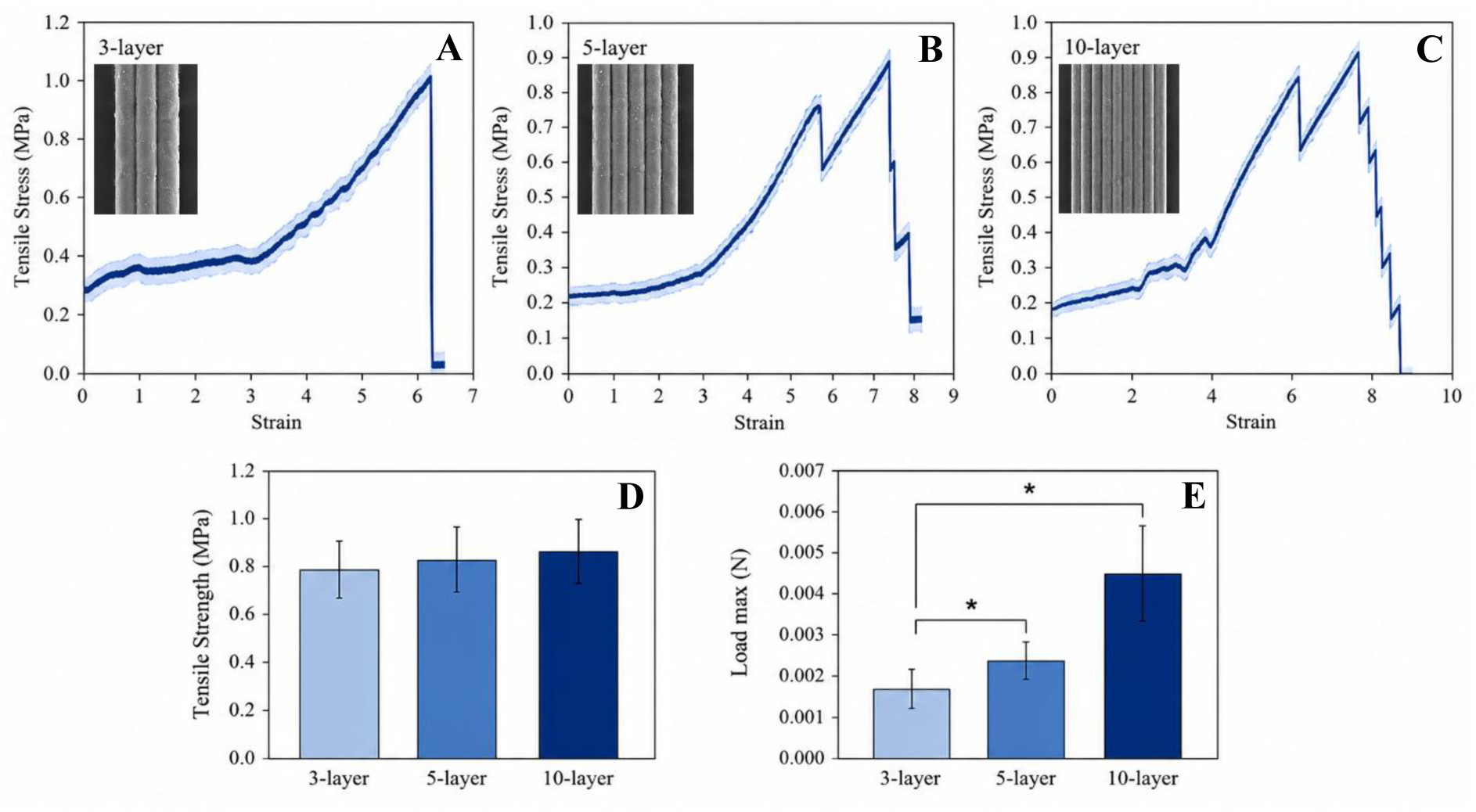
(A–C) Representative stress–strain curves of MEW-fabricated PCL scaffolds with 3, 5, and 10 printed fibre layers, respectively. The strain value of 1 corresponds to 100% elongation. Sequential stress drops indicate the failure of individual fibre layers, while the area under each curve represents the energy absorbed during deformation. The scaffolds exhibit high elongation at break (600–800%), characteristic of their fibrous architecture. (D) Comparison of the uniaxial tensile strength of 3-, 5-, and 10-layer scaffolds. (E) Comparison of the maximum failure load of the corresponding scaffolds. The tensile strength remains nearly constant (∼0.8 MPa), whereas the maximum failure load increases with scaffold thickness due to the larger cross-sectional area. Data are presented as mean ± standard deviation (n = 5).

Reducing the fibre diameter to the microscale improved stress distribution and minimized internal defects, resulting in enhanced deformability without compromising tensile strength. As shown in Figure 1E, increasing the number of stacked fibre layers significantly increased the maximum load-bearing capacity, from approximately ∼2.0 mN for 3 layers to ∼5.0 mN for 10 layers, while the tensile strength remained nearly constant at ∼0.8 MPa. These results indicate that load capacity scales primarily with the number of load-bearing filaments, whereas intrinsic material strength is preserved.

We found constructs with fewer than 10 layers mechanically fragile and difficult to handle reproducibly. Therefore, 10 layers represent the minimum mechanically robust configuration suitable for manipulation and further scaffold assembly. Based on these findings, we selected 10- and 20-layer configurations for all subsequent scaffold-level investigations.

### 3.3. Geometry selection and unit cell optimization: Uniaxial and cyclic tensile testing on MEW scaffolds

Previous work by Iglesias-García et al. demonstrated that rhomboid (diamond-shaped) MEW scaffold architectures provide superior mechanical anisotropy compared to square and rectangular geometries, because the rhomboid geometry supports hinge-like deformation and enhanced functional displacement [15]. The authors established geometry-driven compliance as an effective strategy to direct deformation; however, the analysis was conducted at a fixed scaffold thickness of 10 layers and focussed primarily on effective directional stiffness, without addressing bulk load-bearing capacity or cyclic mechanical behaviour. Moczulska-Heljak et al. investigated the mechanical response of MEW PCL meshes dominated by fibre-scale hierarchy and entanglement, reporting monotonic tensile stiffness, yield force, and yield strain (∼10%), but without consideration of scaffold-level geometry, thickness variation, or anisotropic architectures [25]. It is worth noting that in both studies mechanical behaviour was evaluated exclusively under quasi-static loading conditions, while cyclic tensile performance, which is crucial for the dynamic myocardium, was not assessed.

Consequently, despite advances in geometry-driven anisotropy and process-driven compliance, key questions remain unresolved regarding the mechanical behaviour of multilayer MEW fibres as fundamental load-bearing units, the influence of layer number on scaffold-level strength and deformability, the effect of unit-cell size (porosity) on mechanical integrity, and the cyclic tensile response of MEW-fabricated PCL scaffolds. To address these gaps, we performed a stepwise mechanical investigation, from thickness- and porosity-optimized scaffolds over uniaxial tensile testing to cyclic tensile testing, the latter to evaluate mechanical stability under repeated loading conditions relevant for native cardiac tissue.

To replicate the highly anisotropic mechanical environment of native myocardium, a rhomboid scaffold geometry with a defined aspect ratio of 2 (corresponding to an internal opening angle of 53.1° was selected, based on prior systematic comparisons demonstrating its superiority over square and rectangular architectures in generating directional compliance, through combined experimental mechanical testing and finite element simulations, to produce near-zero effective stiffness along the minor diagonal while maintaining structural integrity along the major axis [15]. Snow et al. performed an extensive mechanical characterization of 10-layer MEW PCL scaffolds with diamond-like architectures, demonstrating that fibre orientation strongly governs tensile stiffness and compliance. Apparent Young’s moduli ranged from 1.8 to 8.0 MPa, depending on fibre angle, with corresponding variations in yield strain and ultimate tensile strength. However, scaffold failure load was not reported, as the highly ductile yielding behavior of MEW PCL and limitations in achievable testing displacement precluded reliable capture of catastrophic failure [26].

We therefore conducted uniaxial tensile tests on precisely cut samples from the scaffold to evaluate scaffold-level mechanical performance in terms of maximum load-bearing capacity. Three scaffold variants were tested: (1) 10-layer R60, (2) 20-layer R60, and (3) 20-layer R120. The 20-layer R120 scaffold exhibited the highest failure load (∼16 N, Figure S3), confirming that increased fibre density and optimized rhomboid geometry substantially enhance structural robustness.

We found that scaffolds with low layer counts and large unit cells generally lacked mechanical integrity and were prone to damage during handling, whereas scaffolds with very small unit cells combined with high layer counts are expected to exhibit increased stiffness, which would limit their ability to deform during cardiomyocyte contraction. Specifically, we found constructs with fewer than 10 layers mechanically fragile and difficult to handle reproducibly. Therefore, 10 layers represent the minimum mechanically robust configuration suitable for further modification and manipulation. For comparison, we also produced 20-layer scaffolds.

Cyclic tensile characterization is essential to evaluate mechanical stability, hysteresis, and strain recovery under repeated loading conditions that more closely resemble the dynamic mechanical environment of native cardiac tissue. We applied cyclic deformations to the 20-layer samples R60 and R120. We used 20 loading cycles (triangular ramps) from 0% to 5% and 10% maximum strain, respectively, at rates of 0.015/s = 1.5%/s (6.5s per cycle) and 0.017/s=1.7%/s (11.50s per cycle), respectively. All experimental values obtained from these tests are summarized in Table S1 and visualized in Figure 2.

**Figure 2.**
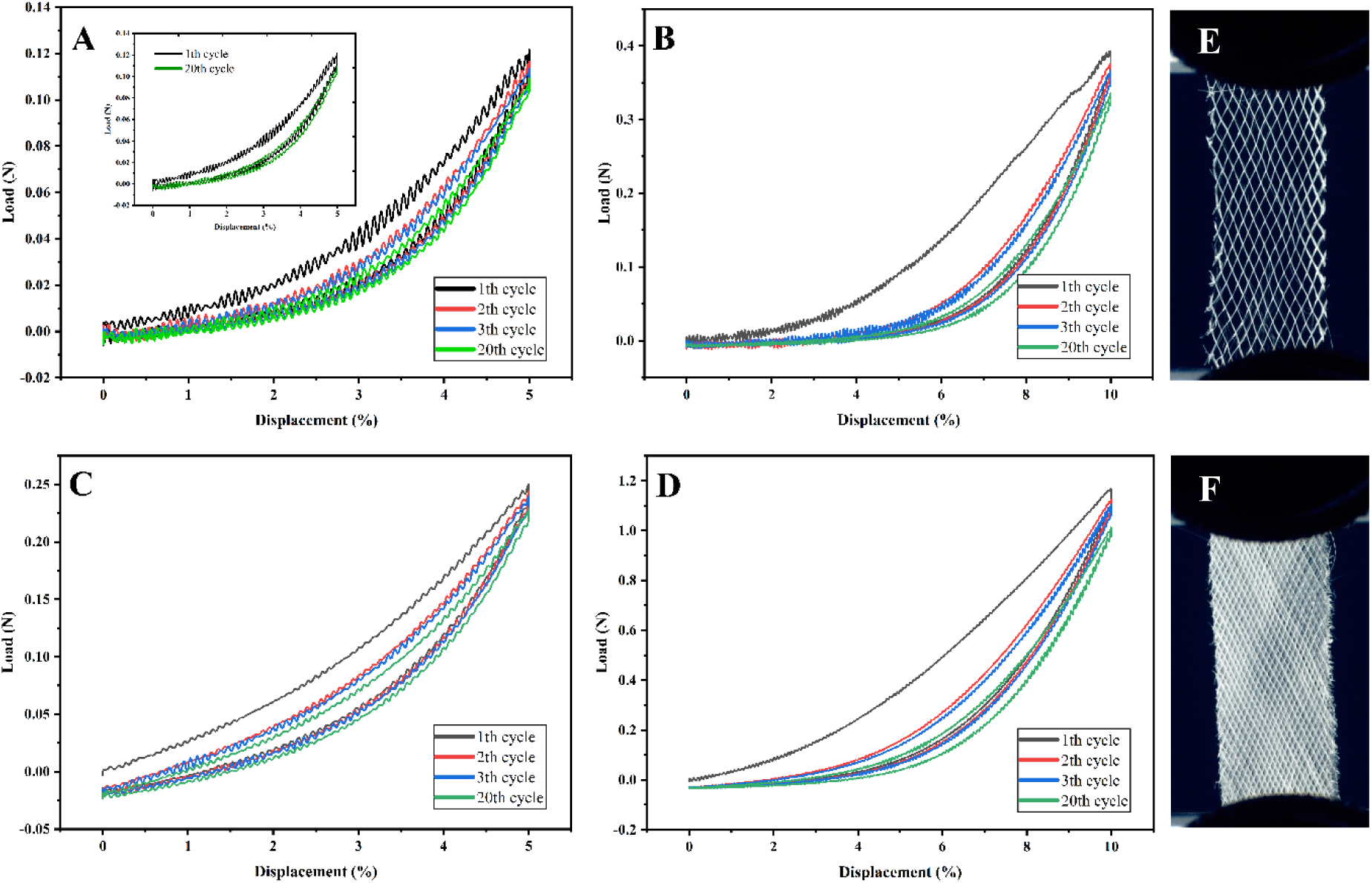
(A) Cyclic mechanical response of 20-layer R60 scaffold under 5% strain. The plot shows the 1st, 2nd, 3rd, and 20th loading–unloading cycles, illustrating progressive mechanical softening. The inset highlights the 1st and 20th cycle. (B) Data for the same scaffold at 10% strain. (C) Data for the 20-layer R120 scaffold under 5% strain. (D) Data for the same scaffold under 10% strain. (E, F) Photographs of 20-layer R60 and R120 scaffolds, respectively, after cyclic loading.

The cyclic tensile behaviour of the 20-layer R60 at 5% strain revealed a relatively soft and elastic mechanical response. Progressive mechanical stabilization was observed, with 26% reduction in energy dissipation, suggesting that internal friction and molecular reorganization occurred predominantly during the early cycles and diminished over time, resulting in a more elastic and stable response under cyclic loading (Figure 2A).

For the same scaffold (20-layer R60) subjected to 10% strain, a higher initial resistance but poorer endurance was observed. Over 20 loading cycles, we found 15% loss in peak strength and 43% reduction in energy dissipation, indicating pronounced internal fatigue and progressive softening (Figure 2B). This substantial loss reduced the scaffold stiffness and elastic recovery capacity, making it less resilient to repeated deformation at higher strain amplitudes.

For the 20-layer R120 scaffold at 5% strain, a stronger and more stable mechanical profile was observed compared to the R60–5% case. Over 20 loading cycles, the scaffold showed an ∼8% loss in peak strength and a 26% reduction in energy dissipation, while maintaining higher absolute load values, indicating enhanced stiffness (Figure 2C). This behaviour reflects a stable elastic response under mild deformation, a desirable feature for load-bearing cardiac patches under mild deformation.

When subjected to 10% cyclic strain, the R120 scaffold exhibited the highest load-bearing capacity among all conditions, as expected for the larger applied strain. Over 20 cycles, it experienced 14% loss in mechanical strength, accompanied by a notable reduction in energy dissipation, indicating a transition toward a more elastic and stabilized mechanical response (Figure 2D). Despite the higher strain, the scaffold maintained substantial load-bearing ability with moderate strength loss, making it a promising candidate for applications requiring both strength and resilience under dynamic conditions.

Overall, the comparison revealed that scaffolds subjected to 5% strain exhibited superior elastic behaviour, characterized by lower energy dissipation and minimal strength degradation, independent of their unit cell size. In contrast, 10% strain led to greater mechanical fatigue, particularly in the R60 configuration, indicating a higher susceptibility to long-term failure. The R120 scaffolds consistently outperformed R60 in terms of strength and stiffness, especially under larger deformation, demonstrating improved structural resilience. These results underscore the inherent trade-off between mechanical robustness and elastic compliance in scaffold design. Owing to its favourable balance between elasticity and mechanical strength, the R120 scaffold was selected for subsequent in vitro studies.

### 3.4. Surface architecture of MEW fibres

Understanding how fibre surface morphology influences cell–material interactions is critical for advancing tissue engineering and biomaterial design. We report various micro- and nanoscale details from MEW PCL, processed under optimized parameters (pressure = 100 mbar, temperature = 80 °C, voltage = 6 kV, and translation speed = 800 mm/min). The high printing precision and structural integrity is visualized in microscale details in ESEM images. Figure 3A presents a 20-layer rhomboid scaffold, featuring a long diagonal of 1 mm and a short diagonal of 0.5 mm, showcasing the geometric complexity achievable through MEW. Figure 3B shows details of three fibres precisely deposited on top of each other, highlighting the fidelity and reproducibility of the MEW process. Figure 3C illustrates the cross-section of a deliberately fractured fibre. The irregular, highly deformed morphology reflects substantial plastic deformation, indicative of the inherent ductility and stretchability of PCL. This plastic fracture profile contrasts sharply with the smooth, brittle surfaces typically seen in more rigid polymers, confirming the mechanical resilience of PCL and its suitability for applications requiring flexibility and robustness.

**Figure 3.**
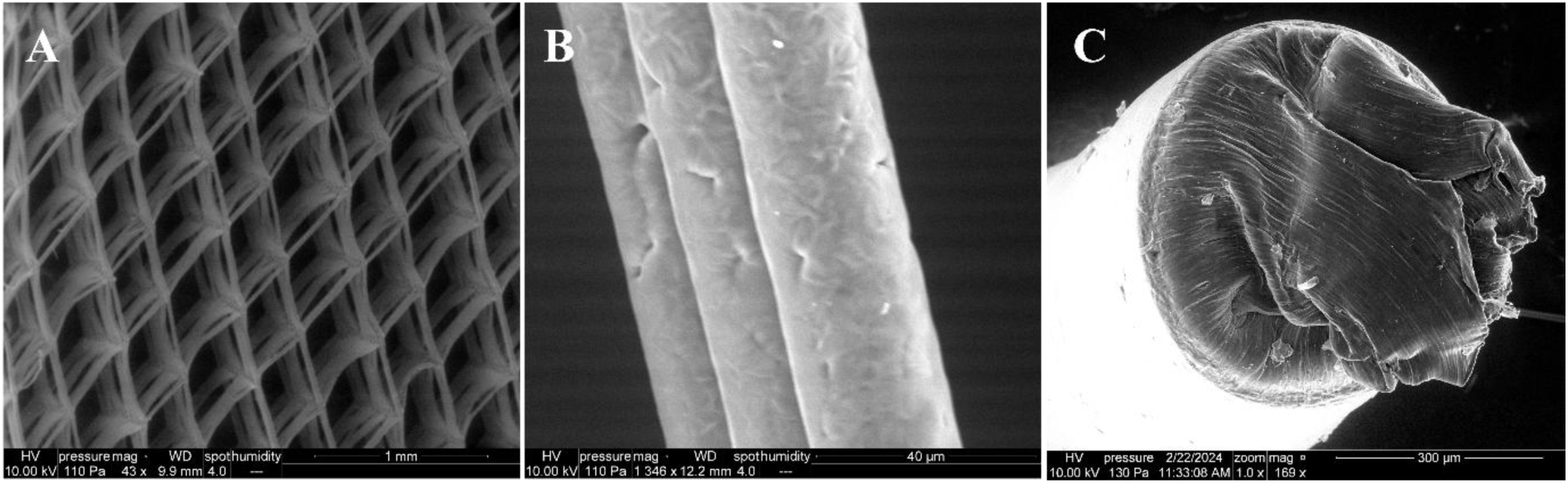
ESEM images of MEW-printed PCL scaffolds: (A) 20-layer rhomboid scaffold (1 mm × 0.5 mm diagonals), (B) three-layer fibre arrangement showing high printing fidelity, and (C) fracture surface of a thick fibre displaying plastic deformation.

Nanoscale surface details such as surface roughness and morphology/topography are detected by AFM. This requires careful placement of the probe tip on top of a single fibre (see Figure 4A). AFM scans acquired over areas ranging from 1 to 5 µm revealed the presence of parallel nanoscale stripes on the fibre surface, visible as partially aligned height variations (Figures 4A and 4B) or as features in the mechanical phase (Figure 4C). The stripe orientation did not consistently coincide with the macroscopic fibre printing direction, similar to the surface features resolved by ESEM (Figure 3B). The height of the stripes was ∼ 3 nm, hence much more than the diameter of a single polymer backbone. We interpret them as highly crystalline domains embedded within the semicrystalline PCL matrix.

**Figure 4.**
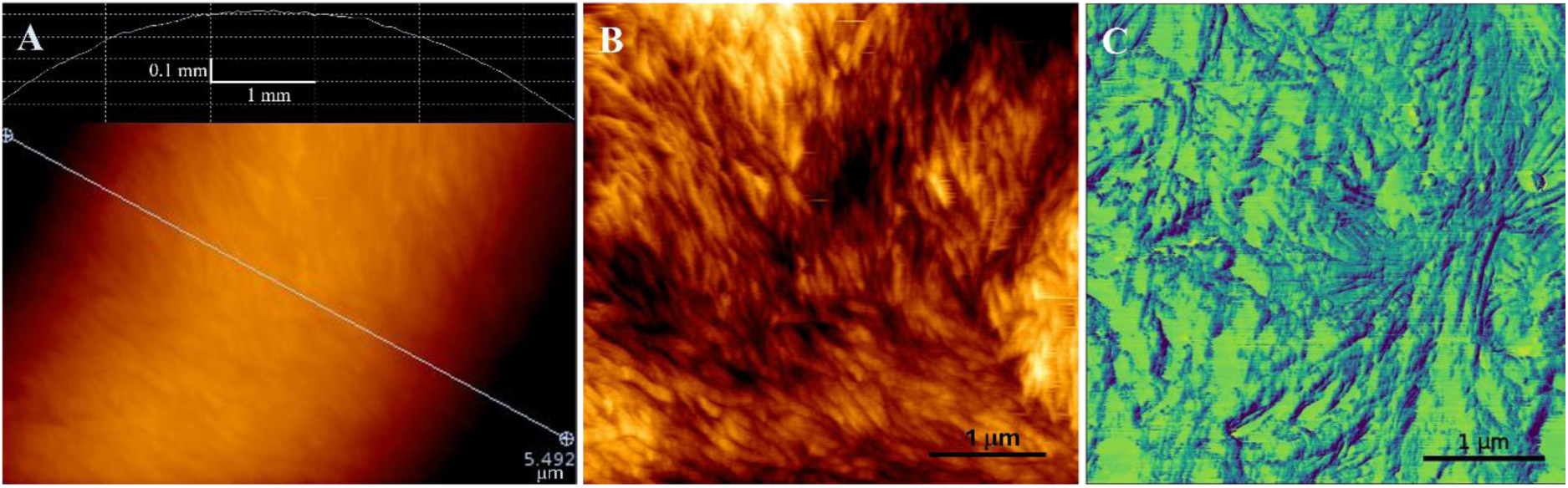
AFM images of three different areas of MEW PCL fibres. (A) Top part of a printed PCL fibre surface. A 5.49 µm long cross-section is marked and shown below the image. (B) In order to visualize details, scans are fitted to a curved baseline. Nanoscale stripes, aligned along variable directions, indicate the presence of crystalline regions. (C) Phase images provide further details, such as a spherulite-type arrangement of stripes in the right middle part.

This observation indicates that MEW not only defines scaffold architecture at the microscale, but also introduces hierarchical structural features at the nanoscale. Such localized crystalline domains are expected to influence local stiffness and surface properties of the fibres, which may in turn affect hydrogel–scaffold interactions and cell mechanosensing in dynamic cardiac constructs. However, further functionalization, as described in the following section, will alter the features – relevant research is out of the scope of this paper.

### 3.5. Incorporation of conductive fillers into bulk PCL

To enhance the electrical performance of MEW-printed PCL scaffolds, a commonly reported strategy is the incorporation of electrically conductive fillers. Given the well-established conductivity of PANI, PPy, and graphene-based materials, these fillers were selected to assess whether bulk modification of PCL could provide a straightforward route to electrically functional MEW scaffolds while maintaining print fidelity.

We first determined the electrical conductivity of the synthesized conductive fillers (see Materials and Methods) with broadband dielectric spectroscopy (BDS) to benchmark their properties. The typical increase of the real part of the conductivity (here Sig’) at high frequencies is not relevant for cardiac tissue, so we focus in the following exclusively on the low-frequency range (≤5 Hz), where all materials exhibited minimal frequency dependence. The PPy and our PANI salts have conductivity values of typical doped semiconductors, 5×10⁻³ S/cm and 5×10⁻⁶ S/cm, respectively (Figure 5A). Among the PANI salts, the sample synthesized using 10% HCl showed slightly higher conductivity compared to those prepared with higher acid concentrations and was therefore selected for composite preparation.

**Figure 5.**
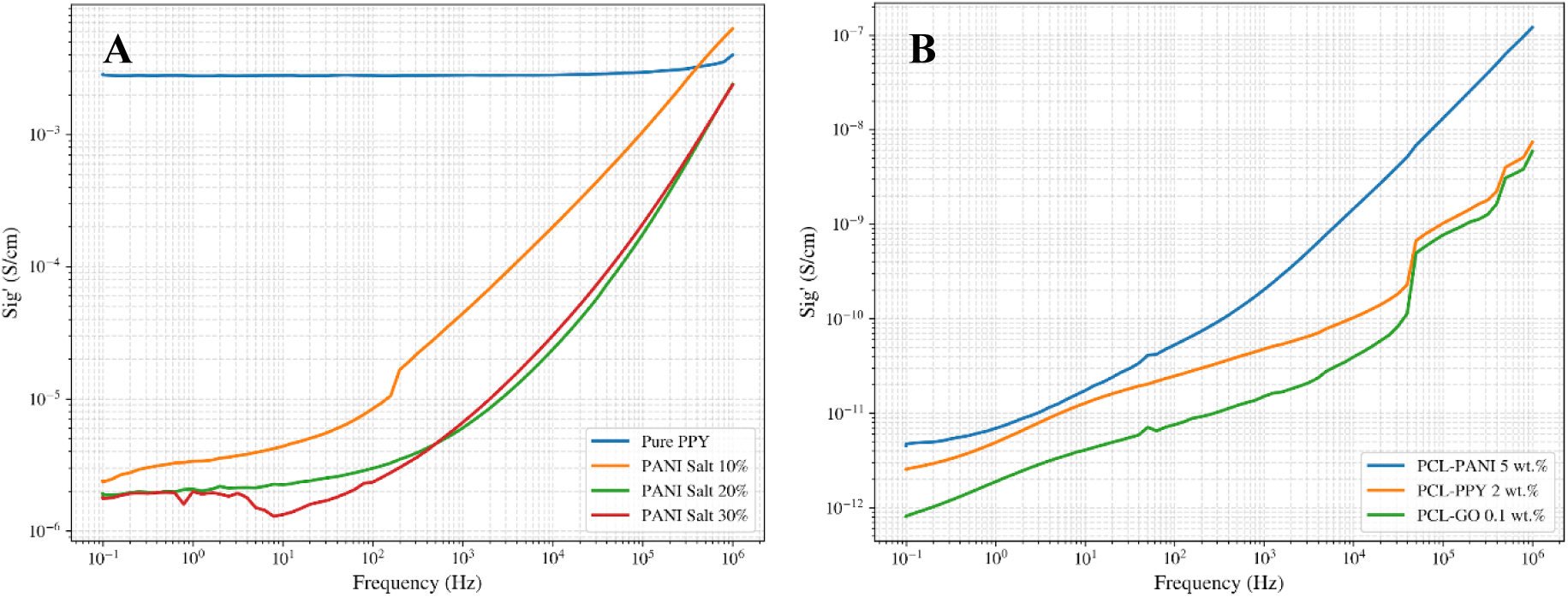
Electrical conductivity analysis with BDS. (A) Conductivity of bulk PPy and of PANI salts synthesized with varying HCl concentrations. (B) Conductivity of PCL-based composites with conductive fillers (PANI 5%, PPy 2%, GO 0.1%).

Composites were formulated by blending PCL with 5 wt.% PANI, 2 wt.% PPy, and 0.1 wt.% graphene oxide (GO), representing the highest filler loadings compatible with stable MEW processing. Increasing the GO content resulted in a pronounced loss of print fidelity (poor melt processability and unstable fibre deposition during MEW), which we also found for the PCL–PANI and PCL–PPy composites. The electrical characterization (Figure 5B) revealed no meaningful enhancement in bulk conductivity over pure PCL (∼10⁻¹⁰ S/cm, not shown), i.e. the composites are insulators. Their conductivities are several orders of magnitude below the physiological range of native cardiac tissue (10⁻³–10⁻⁴ S/cm [27, 28]).

Our interpretation is that, at filler concentrations compatible with MEW processing, the conductive phases do not form a continuous percolated network within the insulating PCL matrix. Hence, we did not try any DC conductivity test (four-probe, see below), but explored surface-based functionalization strategies, which are discussed in the following section.

### 3.6. Gold and PPy coating of MEW PCL

Two independent coating methods were implemented, sputter-coating of (nominally) 20 nm of gold onto the surface of PCL scaffolds (“PCL-Au”) and in situ polymerization of pyrrole on the PCL scaffolds (“PCL-PPy”).

To identify the successful deposition of PPy, we employed Raman spectroscopy of pristine PCL and of PCL-PPy. As expected, PPy lead to additional peaks, of which we highlight the peak at 1572 cm⁻¹ (Figures 6A, B), which we assign to a C=C stretching vibration, which is not present in pure PCL. FTIR analysis (Figures 6C, D) validated these findings: We found a high spectral similarity between the synthesized PPy and a commercial PPy reference in the fingerprint region. E.g., our absorption band at 1283 cm⁻¹ is due to C–N stretching, in agreement with Tabaciarova et al., who reported 1295 cm⁻¹ [29], and our band at ∼1020 cm⁻¹ (aromatic C–H bending) is absent in PCL. Together, the Raman and FTIR results confirm the successful synthesis of PPy on the PCL scaffolds.

**Figure 6.**
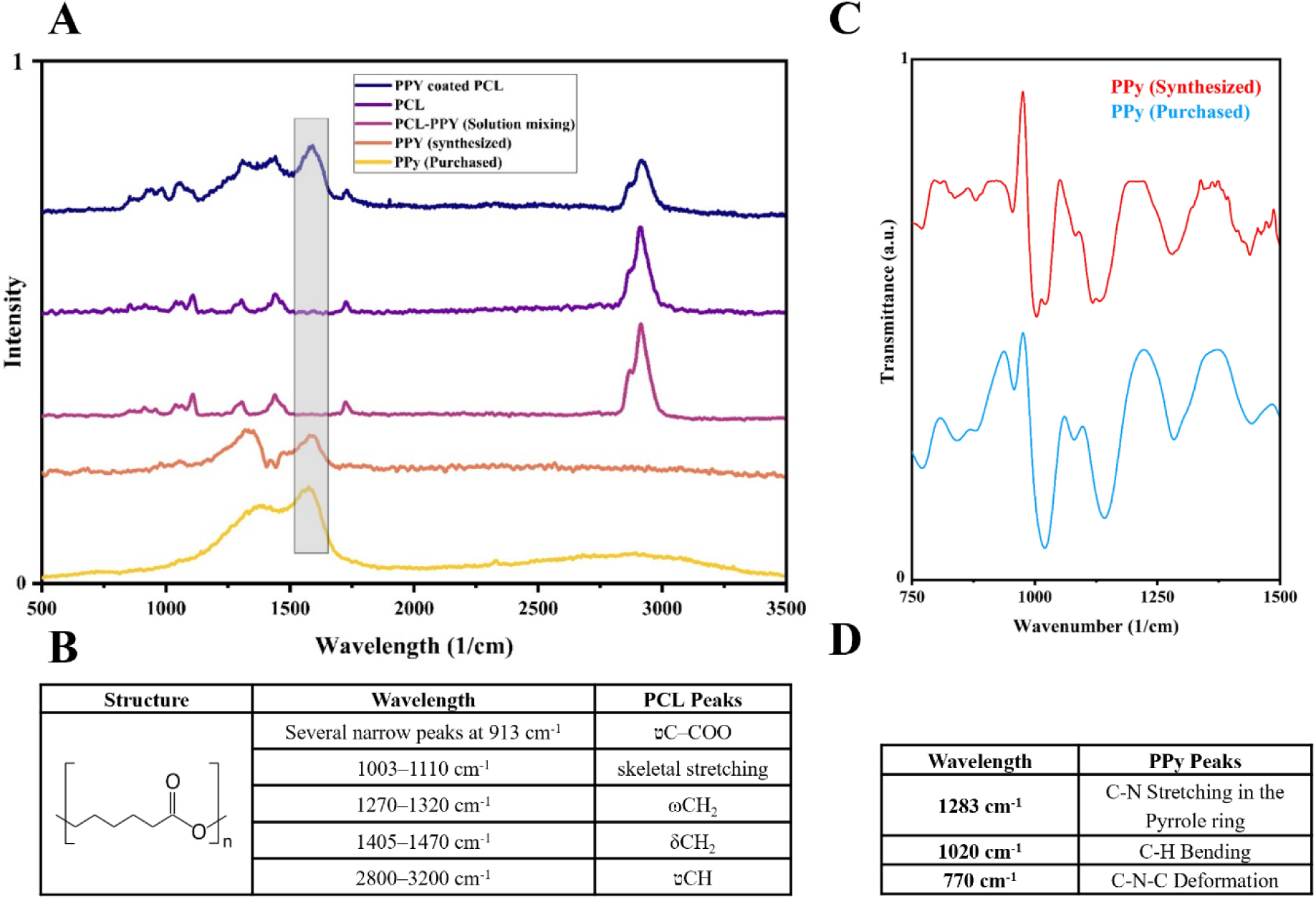
(A) Raman spectra highlighting the PPy-coated PCL scaffold in comparison with pure PCL and PPy reference materials. (B) Peak assignments. (C) FTIR comparison between synthesized and commercial PPy. (D) Peak assignments. Partial peak overlap between PPy and PCL, particularly near ∼1337 cm⁻¹, may influence the clarity of peak assignment in the coated scaffold.

To detect morphological changes caused by PPy, we used optical microscopy and ESEM. Both techniques revealed a characteristic granular surface morphology, showing that PPy deposited as interconnected particles (Figure 7). Crucially, while the coating exhibits a particulate microstructure typical of chemical polymerization, the high density and mutual contact of these particles ensure the formation of a continuous, percolating network capable of electrical conduction, as validated by our subsequent 4-point probe measurements (see below).

**Figure 7.**
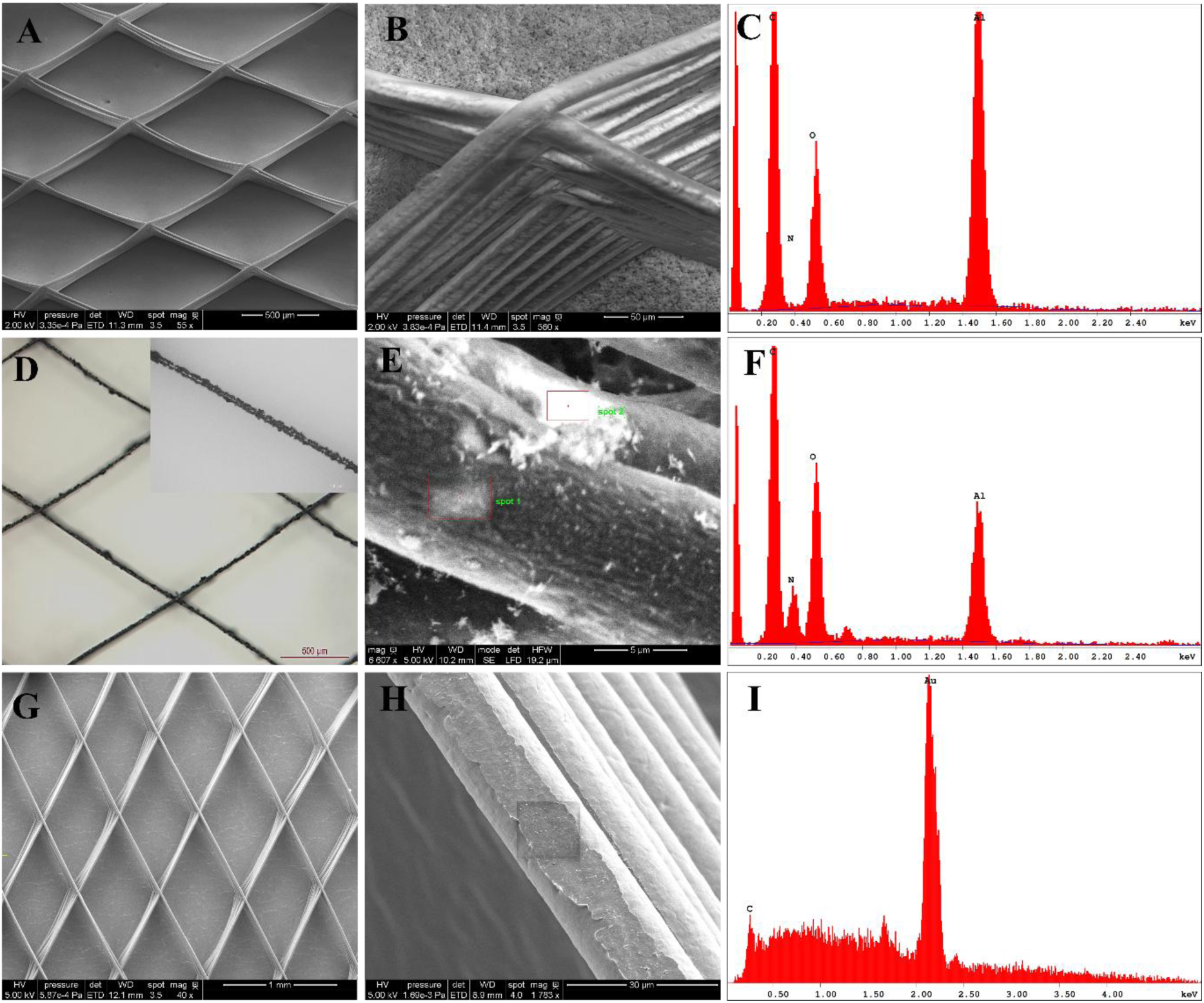
Microscopic and elemental analysis of the scaffolds. (A, B) ESEM images of pure (uncoated) PCL; (D) optical images showing scaffold morphology before and after PPy coating; (E) ESEM image of coated PCL, confirming PPy particle deposition; (G, H) ESEM images of the gold-coated scaffold; (C, F, I) EDS revealing the nitrogen peak in the PPy-coated scaffold (validating the presence of PPy) and the Au peak in the gold-coated scaffold (validating the presence of gold).

The absence of such particles on pure PCL suggests that they are composed of PPy. We validated this hypothesis by ESEM, combined with energy-dispersive X-ray spectroscopy (EDS), conducted at two distinct points on the PCL fibres. EDS results show a low-energy artifact around 0 keV and a signal from the aluminium sample holder at ∼ 1.49 keV (Al K_α_). The carbon (0.27 keV) and oxygen (0.53 keV) peaks, characteristic of the PCL fibre, are present in both spectra. In contrast, the nitrogen peak at ∼0.39 keV is only present for the coated PCL, unequivocally demonstrating the presence of PPy, which contains nitrogen atoms within its molecular structure. (Figures 7C, F). Furthermore, for the gold-coated scaffold, a prominent peak is visible at ∼2.12 keV corresponding to gold (Au M_α_), verifying successful gold deposition (Figure 7I).

In summary, our combined microscale analysis methods confirm the successful polymerization of pyrrole as interconnected particle network on the scaffold surface.

The electrical behavior of the surface-modified scaffolds was evaluated using the 4-point probe method. While both configurations exhibited non-Ohmic, non-linear characteristics (Table 1), the analysis was focused strictly on the lowest current regime due to its physiological relevance. Taking advantage of the uniform gold sputter coating, which yielded a well-controlled thickness of approximately 22 nm, the bulk conductivity (σ = 1/ρ) of this specific variant was calculated. At a physiological current bias of 0.01 µA (corresponding to an R_s_ of 7000 Ω/sq), the gold coating provided an electrical conductivity of approximately 65 S/cm. In contrast, a definitive bulk conversion was intentionally omitted for the PPy-coated scaffolds (R_s_ = 8200 Ω/sq at 0.01 µA). This was done because the in situ polymerization process inherently introduces localized variation in coating thickness across the complex, 3D microfibrous mesh, making any bulk assumption unreliable. Nevertheless, a direct comparison of the raw sheet resistances demonstrates that both surface modification pathways introduce efficient charge transport across the macro-construct. These values represent a substantial improvement over the completely insulating bulk polymer composites discussed earlier. More importantly, at the full patch scale, these low surface resistances establish reliable electrical pathways that satisfy the functional synchronization criteria of native myocardium (10⁻³–10⁻⁴ S/cm [27, 28]) by keeping overall construct resistance to a minimum. Hence, surface coating techniques with gold and PPy successfully enhanced the electrical properties of PCL scaffolds. These two modified scaffolds now serve as promising candidates for subsequent biological evaluations, including in vitro cell culture experiments.

**Table 1.** Comparing the sheet resistance of the fabricated materials using 4-point probe method. The coatings exhibited nonlinear current–voltage characteristics, with higher electrical conductivity observed at increased current levels, particularly in thin gold films where discontinuous domains may form conductive pathways under higher bias.

| Current (μA) | 0.01 | 0.1 | 1 | 10 |
| --- | --- | --- | --- | --- |
| Sheet resistance of the Au coated sample ($\Omega$/sq) | 7000 | 710 | 5.5 | 2.4 |
| Sheet resistance of the PPy coated sample ($\Omega$/sq) | 8200 | 700 | 28 | 106 |

### 3.7. Rhomboidal MEW PCL-Au scaffolds for human contractile cardiac constructs

Once conductive scaffolds were generated, the next step was to produce engineered cardiac constructs that allow us to test if PCL-coating had any biological effect. For the cellular component, hiPSC-CMs were differentiated using chemically defined protocols [16], yielding highly pure cardiomyocyte cultures (Supplemental Video 1). Cardiac constructs were generated by combining hiPSC-CMs (2 million/construct) with rhomboidal scaffolds (PCL, PCL-Au, or PCL-PPy) and embedding the composite in a fibrin hydrogel. The scaffolds consisted of 20 layers, providing sufficient stiffness for handling while preventing limitations in nutrient and oxygen diffusion. Fibrinogen polymerization was induced by thrombin addition (60 min, 37 °C). Constructs were maintained in culture for 2 weeks.

Across the three experimental groups, hiPSC-CMs were densely packed and homogeneously distributed throughout the constructs, populating both regions adjacent to the scaffold fibres and the pore centres (Figure 8A and Figure 9A). Alamar Blue assays demonstrated that control constructs sustained metabolic activity over time, as already reported [15], [30], [31]. Both PCL-Au and PCL-PPy constructs exhibited metabolic activity levels comparable to controls, indicating that the coating strategies did not impair hiPSC-CM metabolism (Figure 8B and 9B).

**Figure 8.**
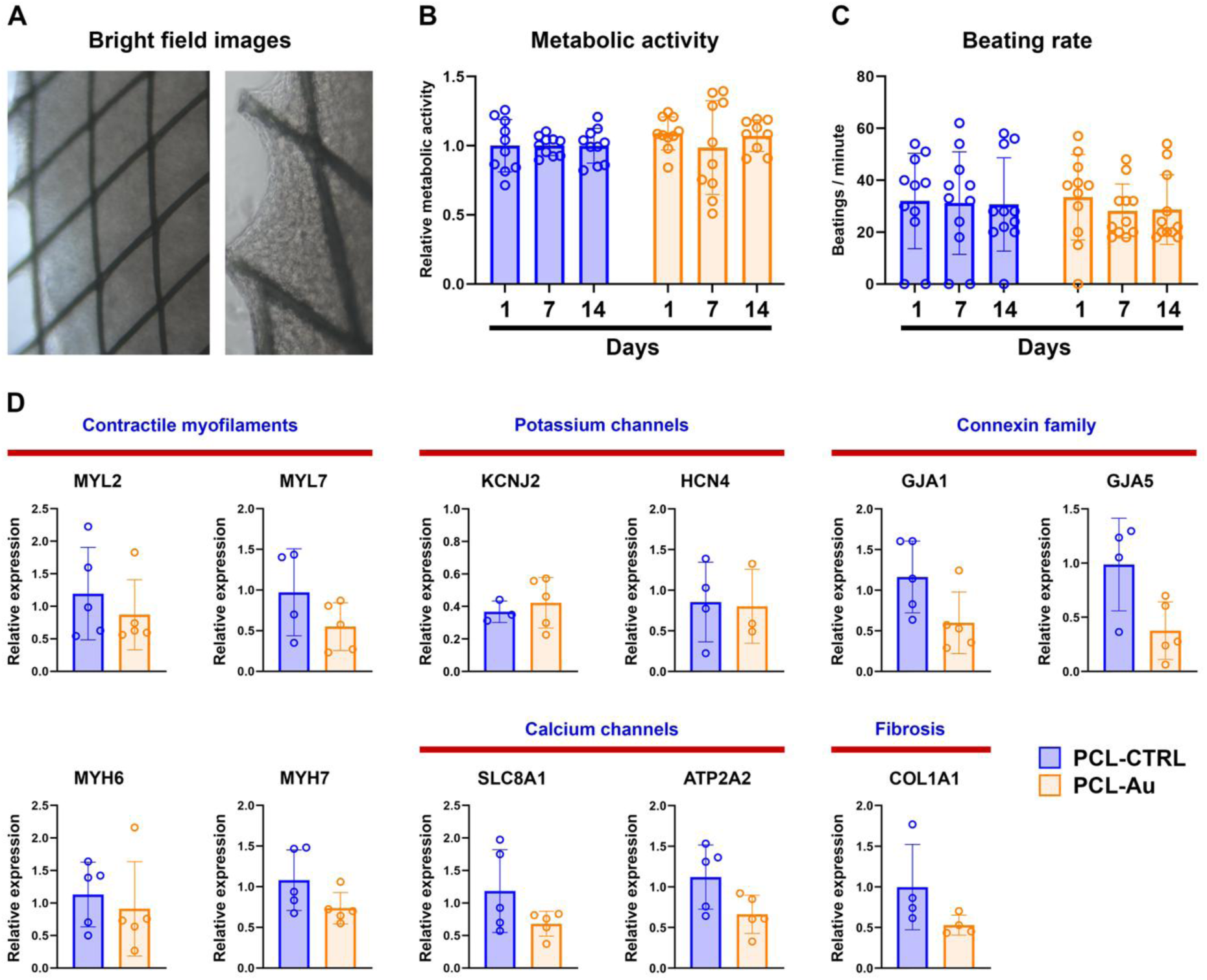
Rhomboidal PCL-Au MEW scaffolds generate functional contractile tissues. A) Representative bright field images reflecting scaffold rhomboidal structure and the homogeneous distribution of hiPSC-CMs along the structure. B) Metabolic activity quantitation employing Alamar blue staining, reflecting metabolic activity maintenance during 14 days tissue culture. C) Evolution of beating rate over time. D) Gene expression analysis by RT-qPCR, relative to PCL-CTRL group, and reflecting a downregulation trend in all cardiac markers. Results are expressed as media with standard deviation. A minimum of 3 independent experiments were conducted.

**Figure 9.**
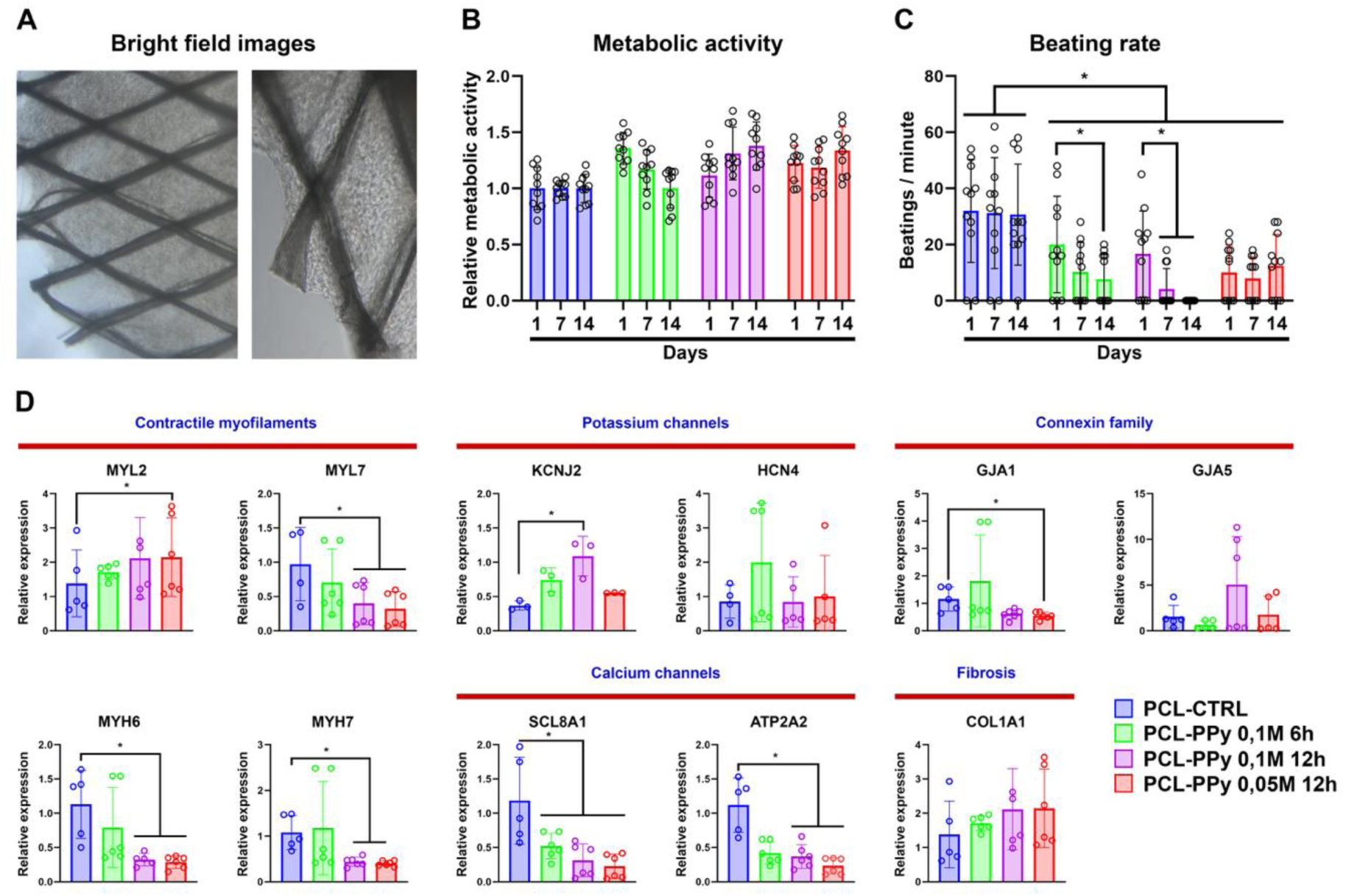
Rhomboidal pore PCL-PPy MEW scaffolds impede construct contraction. A) Representative bright field images reflecting scaffold rhomboidal structure and the homogeneous distribution of hiPSC-CMs along the structure. B) Metabolic activity quantitation employing Alamar blue staining, reflecting metabolic activity maintenance during 14 days tissue culture. C) Evolution of beating rate over time, with a reduced contractile activity over time. D) Gene expression analysis by RT-qPCR, relative to PCL-CTRL samples, reflecting a significant downregulation of cardiac markers in two of the three conditions, and a downregulation trend in the other. Results are expressed as media with standard deviation. *, p < 0,05 by Student’s t-test or Mann-Whitney test. A minimum of 3 independent experiments were conducted.

Regarding functionality, control tissues exhibited spontaneous contractile activity within 24 h of formation, aligned with the minor diagonal of the rhomboidal scaffold. The amplitude of contractions increased progressively, eventually deforming the scaffold. Gold-coated tissues also displayed robust and synchronous beating, with frequencies similar to controls (Figure 8C, Supplemental video 1). In contrast, PPy-coated constructs showed a markedly reduced contractile activity. Their beating rate was significantly lower than that of the control group and further declined over time. Contractile amplitude was severely compromised, with CMs unable to deform the scaffold. Contraction was not observed across the construct, and only localized contractions within the hydrogel were detected (Figure 9C, Supplemental video 2).

Gene expression analysis of PCL-Au scaffolds does not reflect statistically significant differences with the control group. However, a general downregulation trend of cardiac markers in the experimental group is observed. The reduced expression of myosin isoforms (MYL2, MYL7, MYH6, MYH7) suggests an impairment in contractile machinery and a loss of CM maturity, whilst the downregulation of calcium (SLC8A1, ATP2A2) and potassium channels (KCNJ2, HCN4), together with decreased expression of gap junction proteins (GJA1, GJA5), points to compromised electrophysiological function and impaired electrical conduction. Since the cardiac constructs lack cardiac fibroblasts, the reduction in COL1A1 expression suggests a diminished extracellular matrix deposition (Figure 8D). Overall, these findings indicate that PCL-Au scaffolds support the formation of contractile cardiac constructs; however, gold coating does not enhance cardiac functionality compared to pristine PCL and even impairs its physiological development over time.

For PCL-PPy scaffolds, three polymerization conditions were compared. In condition 1 (6 h reaction with 0.1 M pyrrole), transcriptional changes were relatively limited, with only a downregulation of SLC8A1 detected, while the rest of the markers remained comparable to controls. In condition 2 (12 h reaction with 0.1 M pyrrole) and condition 3 (6 h reaction with 0.05 M pyrrole), however, a broad downregulation was observed across most of the analysed genes, including those associated with sarcomeric identity, calcium handling, gap-junction communication, and extracellular matrix deposition. This widespread reduction suggests that prolonged reaction times or suboptimal monomer concentration may compromise scaffold properties in a way that negatively affects CM maturation and functional integration within the constructs.

## Conclusions

In this work, we developed and systematically evaluated electrically conductive MEW-printed PCL scaffolds designed to meet the combined architectural, mechanical, and electrical requirements of engineered cardiac tissue. Through precise optimization of MEW processing parameters, highly reproducible rhomboidal microfibrous scaffolds were fabricated, exhibiting controlled anisotropy and deformation modes that closely resemble those of native myocardium.

Mechanical and electrical performance were found to be strongly governed by scaffold geometry, thickness, and functionalization strategy. Uniaxial tensile profiling of the microfibrous structural units unveiled exceptional stretchability and discrete stress drops, which point towards sequential layer sliding rather than catastrophic fracture. Increasing layer number and reducing unit-cell size significantly enhanced load-bearing capacity and cyclic mechanical stability, while preserving the elastic compliance necessary for cardiac contraction. Crucially, cyclic tensile testing demonstrated that the optimized scaffolds undergo early mechanical stabilization, exhibiting a progressive reduction in hysteresis energy dissipation over consecutive cycles. AFM characterization further indicated that MEW processing induces nanoscale crystalline features on PCL fibres, underscoring the hierarchical nature of the resulting scaffolds and providing additional insight into structure–property relationships across length scales. Attempts to introduce electrical conductivity through bulk incorporation of conductive fillers (PPy, PANI, or GO) were unsuccessful, as these approaches neither achieved physiologically relevant conductivity nor maintained MEW print fidelity. In contrast, surface functionalization via gold sputter coating or in situ PPy polymerization effectively imparted electrical conductivity while preserving scaffold architecture and mechanical integrity, with gold coatings providing superior surface conductivity.

The biological relevance of these design choices was demonstrated using engineered cardiac constructs composed of hiPSC-CMs embedded in fibrin hydrogels. While both conductive coating strategies preserved CM viability, only gold-coated scaffolds supported robust, synchronized, and sustained contractile activity comparable to control tissues. PPy-coated scaffolds, despite being electrically conductive, resulted in mechanically rigid constructs that restricted tissue deformation and impaired functional contraction. Consistent with these observations, gene expression analysis indicated that increased electrical conductivity alone does not promote hiPSC-CM maturation, emphasizing the necessity of balancing conductivity with appropriate mechanical compliance. Collectively, these findings establish that effective cardiac scaffold design relies on the integrated optimization of architecture, mechanics, and electrical functionality, rather than on conductivity as an isolated parameter. This work provides clear design principles and a robust MEW-based platform for the development of next-generation bioengineered cardiac patches with enhanced functional and translational potential.

## Acknowledgements

The authors would like to thank Dr. Javier Aldazabal from Tecnun (Universidad de Navarra, Spain) for his assistance with the cyclic tensile tests, and Dr. Jorge Melillo from the Centro de Física de Materiales (Donostia, Spain) for his support with the Broadband Dielectric Spectroscopy (BDS) measurements.

We acknowledge support by the following grants: Diputación Foral de Gipuzkoa RED I+D “SHIFTE”, MCIN with PID2023-147987OB-C32 (“IVIS-AID”), Lineas Estrateg NEXT “Cardioprint” and the “Maria de Maeztu” Units of Excellence grant CEX2020-001038-M, furthermore the EIC-pathfinder grant 101115292 (“textadna”).

AI was employed for text corrections only.

