## Supplementary Information for "Advancing Cardiac Tissue Engineering: Melt Electrowriting Conductive Polymer-Hydrogel Scaffolds"

### 1. Metadata for DOI Registration

| Item | Title | Description |
|---|---|---|
| Supplementary Figures | MEW Parameters & Mechanics | Parameter optimization, morphological defects, and uniaxial mechanical testing of MEW PCL scaffolds. |
| Table S1 | Cyclic Tensile Performance | Mechanical performance of rhomboid scaffolds under cyclic tensile loading at 5% and 10% strains. |
| Supplemental Video 1 | Au-Scaffold Tissue Beating | Spontaneous and synchronous contraction of hiPSC-CMs on Au-coated MEW PCL scaffolds. |
| Supplemental Video 2 | PPy-Scaffold Tissue Beating | Impaired and localized contractile activity of hiPSC-CMs on PPy-coated MEW PCL scaffolds. |

### 2. Supplementary Figures

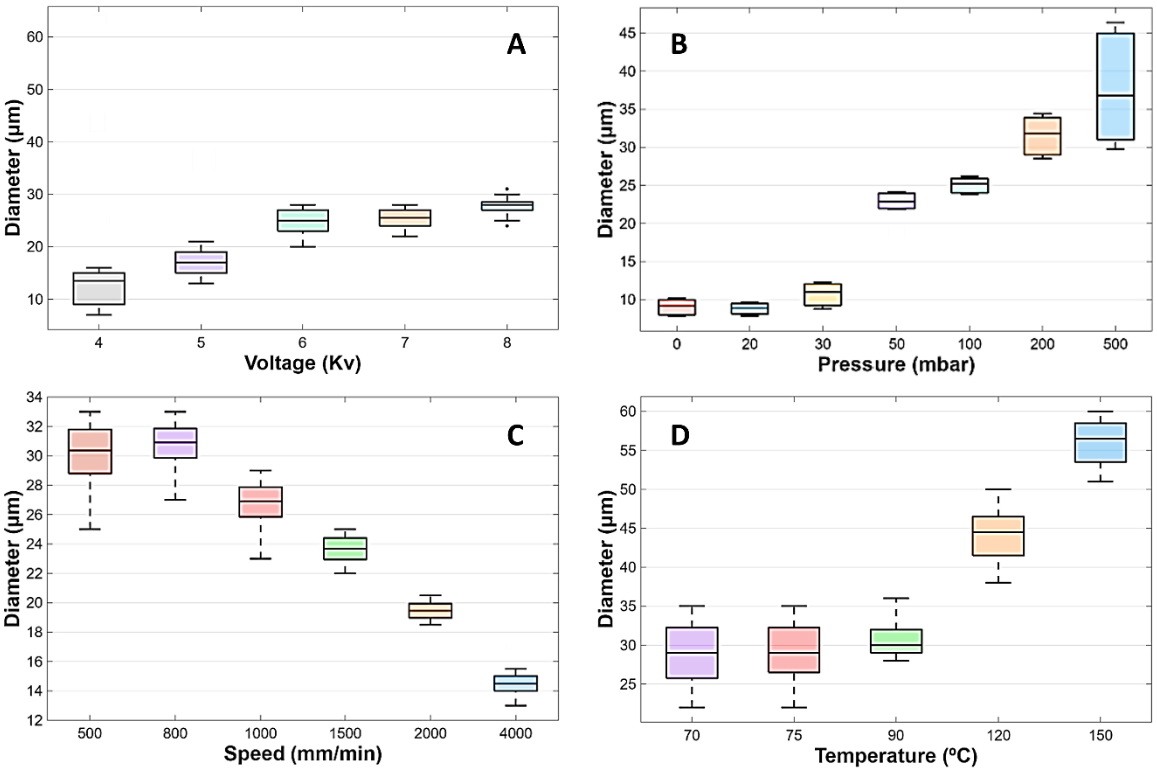


***Figure S1.*** *Effect of key MEW processing parameters on fiber morphology and diameter. (A) Voltage: Stable fiber formation occurs between -6 and -7 kV, with inconsistent printing at lower voltages and fiber splitting at -8 kV. (B) Pressure: Fiber diameter increases significantly with pressure, from 8.82 µm at 20 mbar to 37.70 µm at 500 mbar. (C) Printing speed: Increasing speed from 500 to 4000 mm/min reduces fiber diameter by over 50%, though excessive speed affects trajectory control and printing fidelity. (D) Temperature: 70–90 °C supports suspended fibers; >90 °C reduces suspension, >150 °C causes buckling. Diameter grows from ~29.15 µm (70 °C)*

*to ~45.45 µm (120 °C). Each data point represents the mean diameter from five independent samples, with each sample measured at five different locations. Error bars indicate the standard deviation reflecting both local heterogeneity and sample-to-sample variation.*

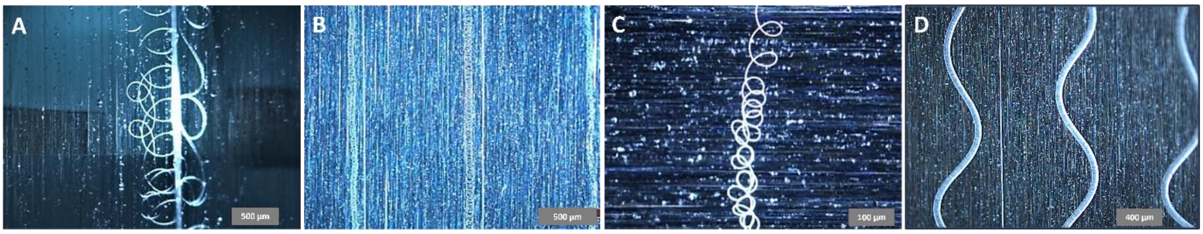


***Figure S2.*** *Morphological irregularities under suboptimal MEW conditions. (a) Fiber splitting at -8 kV: simultaneous formation of a thick straight fiber and a thin curly fiber. (b, c) Thin, curly fibers formed at very low pressure (0–20 mbar), with diameters in the submicron range. (d) Sinusoidal fiber patterns observed at high melt temperature (150 °C), indicating jet instability due to reduced viscosity.*

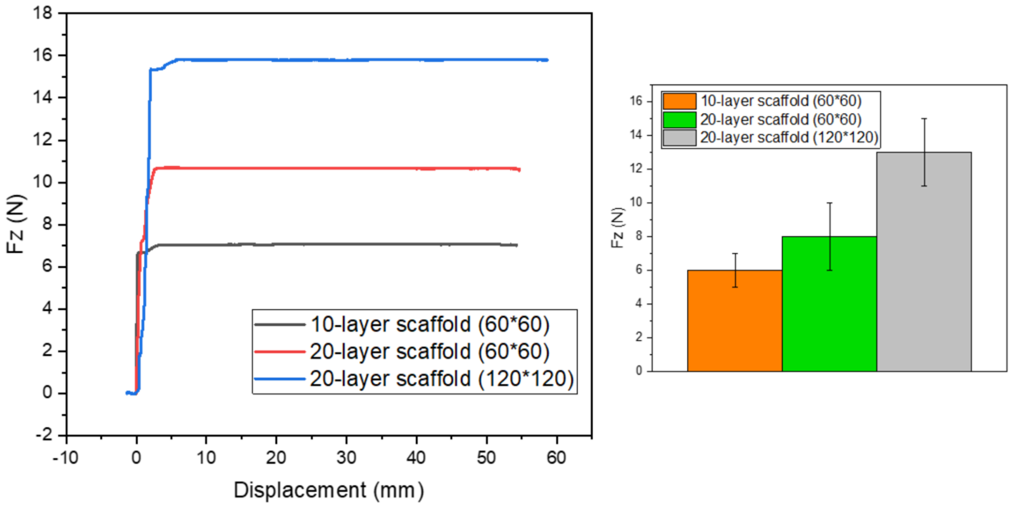


***Figure S3.*** *Uniaxial tensile properties of MEW-fabricated rhomboid PCL scaffolds with varying layer counts (10 or 20) and unit cell densities (60×60 or 120×120). Increasing fiber density and layer number enhanced the maximum failure load, with the 20-layer, 120×120 configuration (Rhomb-120) achieving the highest load (~16 N) while maintaining sufficient flexibility for cardiac tissue engineering applications. Data are averaged over five independent samples, and error bars show the standard deviation corresponding to sample-to-sample variability.*

### 3. Supplementary Tables

***Table S1.*** *Comparative Mechanical Performance of Rhomboid Scaffolds with Different Unit Cell Sizes and Strain Levels Under Cyclic Tensile Loading. Maximum load values under 10% strain are inherently higher due to increased deformation. These values reflect the scaffold's response to the applied strain and should not be interpreted as an absolute measure of intrinsic mechanical superiority. Instead, fatigue performance, strength retention, and hysteresis reduction should be considered together for a comprehensive evaluation.*

| Scaffold Type | Max Load (1st Cycle), (N) | Max Load (20th Cycle), (N) | Strength Loss (%) | Hysteresis Area (1st Cycle), (%.N) | Hysteresis Area (20th Cycle) |
|---|---|---|---|---|---|
| R60-5% | 0.120 | 0.110 | 8.3 | 0.164 | 0.122 |
| R60-10% | 0.390 | 0.331 | 15 | 0.954 | 0.545 |
| R120-5% | 0.249 | 0.229 | 8 | 0.384 | 0.286 |
| R120-10% | 1.167 | 1.007 | 13.7 | 3.338 | 2.063 |

### 4. Supplemental Video Legends

***Supplemental Video 1.*** *Spontaneous and synchronous contraction of hiPSC-CMs on Au-coated MEW scaffold. Video showing the spontaneous contractile activity of hiPSC-CMs integrated with a 20-layer Au-coated rhomboidal MEW PCL scaffold within a fibrin hydrogel. The tissue demonstrates robust, high-amplitude, and synchronous beating aligned with the minor diagonal of the scaffold structure.*

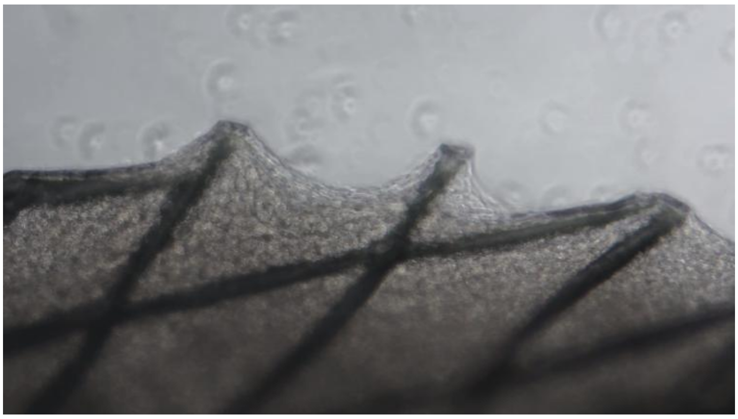

***Supplemental Video 2.*** *Compromised contractile activity of hiPSC-CMs on PPy-coated MEW scaffold. Video depicting the contractile behavior of hiPSC-CMs on a PPy-coated rhomboidal MEW PCL scaffold. The construct exhibits markedly reduced beating frequency and severely compromised contraction amplitude, with localized contractions failing to deform the scaffold construct globally.*

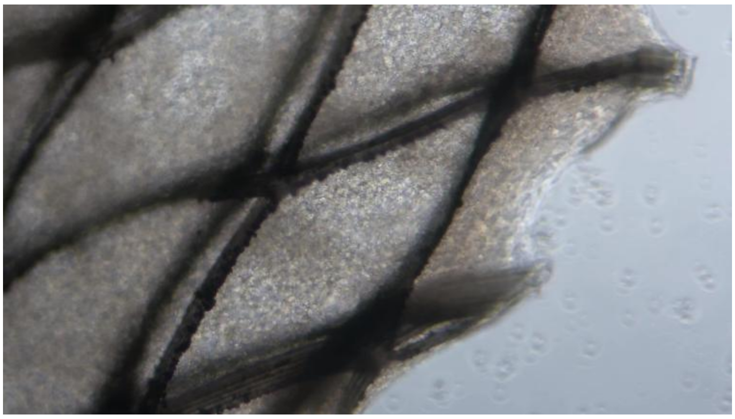
